# Interceptive capturing in large-billed crows: Velocity-dependent weighing of prediction of future target location and visual feedback of current target location

**DOI:** 10.1101/2020.05.06.080671

**Authors:** Yusuke Ujihara, Hiroshi Matsui, Ei-Ichi Izawa

## Abstract

Interception of a moving target is a fundamental behaviour of predators and requires tight coupling between the sensory and motor systems. In the literature of foraging studies, feedback mechanisms based on current target position are frequently reported. However, there have also been recent reports of animals employing feedforward mechanisms, in which prediction of future target location plays an important role. In nature, coordination of these two mechanisms may contribute to intercepting evasive prey. However, how animals weigh these two mechanisms remain poorly understood. Here, we conducted a behavioural experiment involving crows (which show flexible sensorimotor coordination in various domains) capturing a moving target. We changed the velocity of the target to examine how the crows utilised prediction of the target location. The analysis of moment-to-moment head movements and computational simulations revealed that the crows used prediction of future target location when the target velocity was high. In contrast, their interception depended on the current momentary position of the target when the target velocity was slow. These results suggest that crows successfully intercept targets by weighing predictive and visual feedback mechanisms, depending on the target velocity.

## Introduction

Capturing a moving target (interception) is a fundamental behaviour of predators and requires tight coupling between the sensory and motor systems. Two major behavioural mechanisms of interception have been proposed: pure pursuit and proportional navigation (1, 2; also see the Methods section below). Pure pursuit is a behavioural mechanism in which the predator chases a moving target by heading directly to the location of the target. With this mechanism it is assumed that the predator minimises the angles between its own direction of movement and the relative position of the target. Involvement of the pure pursuit mechanism in interception has been suggested in various predatory and non-predatory species [bees (3), flies (4, 5), beetles (6), bats (7, 8)].

In another mechanism of interception, proportional navigation, it is assumed that animals determine their direction of movement according to the angular displacement elicited by movement of the target. Under ideal conditions, animals keep their angular position constant relative to the target. Proportional navigation is employed by various aerial, aquatic, and terrestrial species [dragonflies (9, 10), bats (11, 12), hawks (13), flies (2, 14), fish (15), and beetles (16)]. Proportional navigation has the advantage of simplicity of implementation, since it does not require allocentric coordinates of self and target location (2, 17). Rather, only retinal coordinates are sufficient—if the animal determines its direction of movement in order to keep the retinal position of the target constant.

In both pure pursuit and proportional navigation, it is presumed that the animal utilises visual feedback from the current target location. However, several studies have indicated that not only the current location but also the predicted future location of the target have important roles in successful capture of a moving target [dragonfly (18), humans (19), macaque monkey (20), pigeon (21, 22), snake (23), zebrafish (24)]. Mischiati and colleagues (18) tracked aerial predation of dragonflies (*Plathemis lydia*) and showed that the dragonflies predictively oriented their direction of movement, suggesting formation of an internal model of the movement of the prey (18). In humans, the motion of reaching for a moving target is also based on prediction of target location (19). It is evident that feedforward, prediction-based and visual feedback-based motor control are not mutually exclusive mechanisms. In fact, reaching movement in primates is controlled by both feedforward (predictive) and feedback mechanisms based on vision (25, 26). Given that the movement speed of prey varies in natural situations, the predator may be required to weight the location prediction (feedforward) and visual cues (feedback) according to the speed of the target.

Corvid birds are suitable subjects for the investigation of such sensorimotor control mechanisms, given that their sensorimotor flexibility has been reported in various domains, including motor learning (27, 28), feeding behaviour (29), tool-using (30, 31, 32), and physical reasoning (33). The feeding behaviour of birds, ‘pecking’, is a behaviour analogous to arm-reaching of primates, and have been investigated mainly in pigeons (34). Pecking behaviour in pigeons has been reported under control mechanisms different from those exhibited in arm-reaching of primates. Unlike primate reaching, pecking has been regarded as a feedforward movement in which visual feedback has little effect during movements (29, 34-36). In fact, interception of moving targets was investigated in pigeons, and it was shown that pigeons used prediction of future target location, but not visual guidance after initiation of movements (21, 22). In contrast to the behaviour of pigeons, the pecking behaviour of crows is controlled by visual feedback (27, 28). Thus, investigation of target interception by crows would provide ample opportunities to examine the weighing of feedforward (prediction) mechanism and the visual feedback mechanism.

In the present study, we aimed to resolve three related questions; first, what behavioural mechanism would crows adopt to intercept a moving target; second, whether crows would use prediction of future target location to capture a moving target; and third, if crows depend on prediction of future target location, whether weighting of the prediction would vary according to the speed of the moving target. It was predicted that crows would greatly weigh the predictive mechanism for a fast-moving target, whereas for a slower moving target, they would determine their movement depending on the current target location. To answer these questions, we performed a behavioural experiment in which crows intercepted a moving target (food). The moment-to-moment head movements of the crows were analysed with mathematical models. Additionally, computational simulation of models was performed to validate whether the models capture important aspects of the movements, including the rate of successful capture and movement dynamics.

## Materials and Methods

### Animals and housing

Three large-billed crows (*Corvus macrorhynchos*, two males and one unknown sex, body mass: 520–810 g) were used in the study. All birds were experimentally naive and wild-caught in Tokyo, as authorised by the Environmental Bureau of Tokyo Metropolitan Government (Permission #4005). Crows were housed individually in stainless steel-mesh home cages (43 × 60 × 50 cm, width × depth × height) for approximately 2 weeks for the experimental periods plus three days for acclimation to the experimental chamber. Individuals were placed side-by-side to allow visual and audio-vocal social communication with one another. During the experimental period, crows were regularly transferred into an outdoor aviary (1.5 × 2.8 × 1.7 m, width × depth × height) for bathing and direct social interactions with other conspecifics. At the end of the experimental period, the birds were transferred back to a larger outdoor aviary for group housing (100 m^2^ × 3 m in height) and used for other behavioural studies. Feeding was controlled during the experimental period: specifically, no food was provided for 3 hours before the daily session, but sufficient food was given after the session in the home cage. Daily diets contained dry foods, cheese, and eggs. Water was freely available in their home cage. The room was maintained at 21 ± 2 °C in a 13 L:11 D cycle with light onset at 08:00. The experimental and housing protocols adhered to Japanese National Regulations for Animal Welfare and were approved by the Animal Care and Use Committee of Keio University (no. 14077).

### Apparatus

The experiment was performed in a chamber (63 × 60 × 45 cm, width × depth × height) separated by transparent acrylic plates into two rooms: one for a crow (60 × 45 cm, width × depth) and the other for the target presentation apparatus (Fig. 1). Acrylic plates were set side by side with an 8 cm slit to allow the crow to approach a moving target. The target was presented with reciprocating linear motion using a custom-made apparatus produced from a 3-D printer. The apparatus consisted of a motor-driven timing belt, which was controlled by an Arduino board and custom-made software. As a pecking target, a sphere-cut piece of cheese (2 × 1 × 0.5 cm, width × depth × height, 0.8 g) was attached to the tip of a metal wire set on the timing belt (approximately 20 cm above the floor). A high-speed camcorder (300 frames/s; Gig-E 200, Library Inc., Tokyo, Japan) was located above the chamber (150 cm).

**Figure 1:**
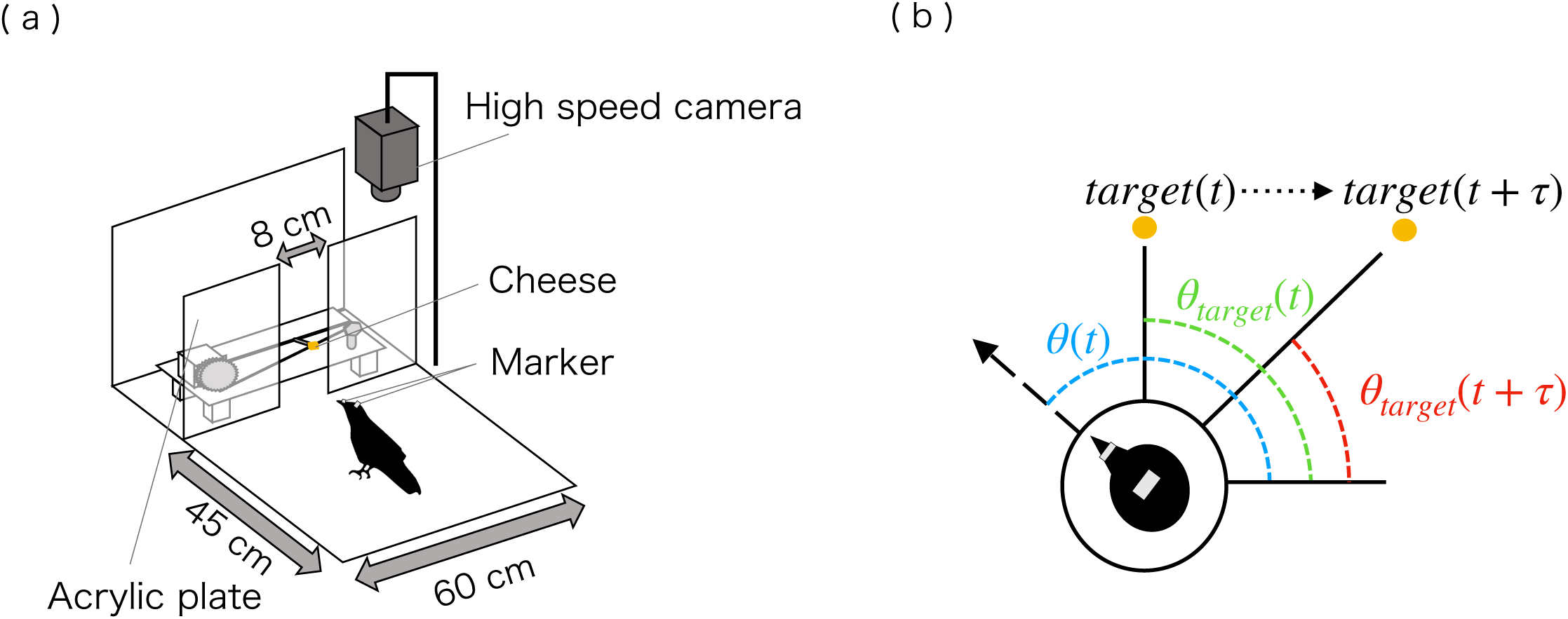
(a): Schematic diagram of experimental chamber. A sphere-cut small piece of cheese as an interception target was moved along a belt. The crow was allowed to access to the target though an 8-cm slit. Beak and head movements were recorded with a high-speed camcorder mounted above the chamber. (b): Definition of angular deviation used in modelling. Head orientation, *θ*(t), was calculated as the angle between the beak-head and an arbitrary fixed reference vector. The angular target position, *θ* _*target*_ (t) was similarly defined as the angle between the head-target and reference vectors. The parameter, *τ* represents future target location and is used in the modelling to determine whether the crows adopted predictive movement in their interception behaviour.

### Procedure

The experiment consisted of three target velocity conditions: 9, 15, and 30 cm/s. Before the start of a trial, the target began to move with a constant velocity while the slit was closed with a transparent acrylic plate. At the start of the trial, the acrylic plate was pulled up by the experimenter, allowing the crow to intercept the moving food. The trial was stopped at the point of the first successful or failed capture or when 2.5 min. had elapsed from the start of the trial without any capturing behaviour. The slit was again closed immediately after the end of the trial. After an inter-trial interval of 0.5–1.5 min., the next trial was started. A single session consisted of 12 successive trials. Within a single session, target velocity conditions were changed every third trial. Thus, each session contained all of the target velocity conditions. Two sessions were conducted per individual in a day, with a 1–2 h interval between sessions. For each crow, a total of six sessions (with 1–2 days interval between every two sessions) were conducted. The order of conditions in all sessions was counter-balanced to offset the order effect.

### Trajectory extraction

The horizontal movement of intercepting a moving target was video captured. Removable square-cut white markers (1 × 1 cm, width × depth) were attached to the head of the crow and the middle of the upper beak. The trials in which the head of the crow passed through the slit were omitted from the trajectory extraction and analysis, since in such trials the crow did not intercept the target but waited for it without overt horizontal head movement. The trajectories of the head, beak, and target were extracted using tracking software (Move-tr/2D v. 7.0, Library Inc., Tokyo, Japan). Extracted x-y coordinates were smoothed using a smoothing splines function to remove tracking noise. Specifically, we applied generalised cross-validation in order to set a smoothing parameter with minimal generalisation error. Employing these trajectories, we calculated moment-to-moment head orientation relative to the target position. These physical quantities were employed for the mathematical models below.

### Mathematical models

Our models were modified versions of two types of foraging models, used in previous foraging studies: the pure pursuit model and proportional navigation model, as described by Fabian et al. (2) In ordinary pure pursuit and proportional navigation models, movements of head orientation are expressed as follows;

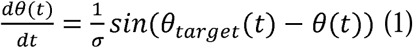

where *α* represents a positive weighting parameter, *θ* _*target*_ denotes the angular position of the target, and *θ* represents the head orientation. The pure pursuit model assumes that animals determine their movement orientation according to angular deviation from the target location (i.e.,

*θ* _*target*_ – *θ*). In contrast, in proportional navigation, the angular movements of head orientation are proportional to the angular velocity of the target position (as noted by Fabian et al. (2)).

In both models it is assumed that animals determine their movement orientation based on the current target location; in other words, visual feedback of current target location is used in a mechanism of interceptive behaviour. However, it is probable that animals adopted not only a feedback mechanism but also a feedforward mechanism, predicting the future target location, especially in the case of fast-moving targets. To examine the contribution of the two mechanisms to the foraging behaviour, we added a prediction term in each model as follows:

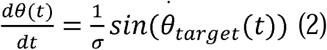

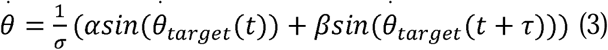

where *τ* is a positive parameter for prediction, which indicates the predicted distance of the target from its present location, *α* and *β* are the parameters of the ratio, which determine the weighting of current and future target location. If *τ*= 0, these models are reduced to models (1) and (2). To investigate the ratio of the contribution between current location of the target and predicted future location, parameters *α* and *β* were constrained as follows:

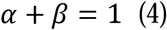

To achieve this constraint, parameters *α* and *β* are defined as *α* = sin^2^(*φ*), and *β*= cos^2^(*φ*), and a hyperparameter *φ* controls the ratio of the contribution of current and future (predicted) location of the target.

### Parameter estimation

Our models include three parameters: *τ*, φ, and *σ*. We estimated these parameters using the Bayesian inference. Because parameter *τ* is a discrete value in the real data (depending on the video flame rate), it caused a convergence problem in the estimation of the joint posterior distribution of all three parameters. Thus, we calculated and compared marginal likelihood of the models with each parameter *τ*. Employing the highest likelihood *τ*, we estimated other parameters. For the other two parameters, Bayesian statistical models were defined as follows:

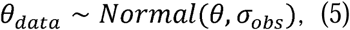

where, *θ*_*data*_ is the angular velocity of the head calculated from experimental data, *θ* is model-generated angular velocity, expressed in equation (2) or (3). *α*_*obs*_ is a parameter of Gaussian distribution, which is estimated in each model. We defined prior distributions of each parameter as follows:

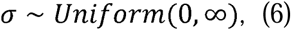

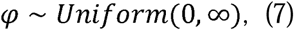

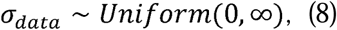

We set a single highest likelihood *τ* for each model and estimated *φ* and *σ* for every condition. These calculations and estimations were conducted using MCMC sampling with four Markov chains of 1,800 iterations, 1,500 warmup, and thinning interval of 3. The convergence of models was checked by Gelman-Rubin statistics 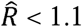 and visual inspection of MCMC traceplots.

### Model comparison

To compare the two interception models, we used Bayes factor as an indicator of the relative explanatory power of the two models. Bayes factor is a way to compare models by quantifying the support for a model over another (37). We used a prior distribution and a posterior distribution of a model with highest likelihood *τ* in each model to calculate the Bayes factor. The calculation was conducted using the bridge sampling method. In addition, the coefficient of determination was calculated to examine the absolute fitting accuracy. Regarding the coefficient of determination, we used the mean of the posterior predictive distribution of angular velocity and observed the values from the experiment.

### Computational simulations

To examine which parameter set had a high probability of generating successful capture, we conducted computational simulations of interceptive behaviour. Specifically, we simulated the moment-to-moment interceptive movement of the crows by assuming that the simulated crows changed their head direction according to the pure pursuit model or proportional navigation model described in the differential equations (3) and (4) in every time step. For simplicity, the translational moving velocity of a simulated crow was fixed at 30 cm/s based on the experimental data. To mimic the experimental situation in the simulation, the initial position of the target was randomly generated along the target path in every trial. The initial relative position of the simulated crow to the target and its beak direction were sampled from the empirical distributions of the experimental data, which were created from the resampling of experimental data using the smooth bootstrap method. A trial was stopped when the crow’s beak reached the target or path of the target movement. Three thousand trials were performed for each parameter set of *φ* and *τ*, ranging from 0 to *ττ*/2 by 63 steps and 0 to 0.2 s by 61 steps, respectively. For all simulations, the parameter *σ* was fixed to prevent the angular velocity of the simulated crow from exceeding the values in the experimental data.

A simulated trial was regarded as successful when the distance between the beak of the simulated crow and the target was less than 2 cm at the end of a trial, judging from the width of the beak and the target. Trials that were not terminated at the centre (8 cm width) of the target path were excluded from the success rate calculation, because such a movement would not occur in an actual experiment. Trials in which the initial angular deviation of the crow was larger than |0.4| (rad) were also excluded from the calculation of the success rate, because these parameters could not feasibly generate successful capture of the target. This would be valid operation, since the real animal often starts pursuit with little angle deviation by using head saccades before movement initiation (13). However, our aim was not to examine such a preparatory response, but rather the interceptive movement itself. Thus, we constrained the initial angular deviation to values smaller than |0.4| (rad).

Additionally, to compare movement trajectory of the simulated and real crows, the angular deviation of trajectories was calculated (θ(t) in Fig. 1b; also see Fig. 5a). Because real crows orient their beak beyond the direction of target movement (‘overshooting’ their direction; Fig. 5a), we considered it worth examining whether the simulation could capture this movement feature. Each simulated trial was calculated using the fourth-order Runge-Kutta method. All statistical analyses were performed in an R v3.6.1 environment (38). Bayesian modelling was built through Stan modelling language (39), and the simulation was performed using Julia v1.2.0 (40).

## Results

A total of 179 responses were successfully recorded and analysed (54, 62, and 63 from the 9, 15, and 30 cm/s conditions, respectively). The 95% credible intervals (CI) of the weighting parameter ϕ varied depending on the target condition in both foraging models (Fig. 2a). The 95% CI of 9, 15, and 30 cm in the proportional navigation model were [1.23 1.53], [1.32 1.56], and [0.68 0.81], and those in the pure pursuit model were [1.07 1.48], [0.84 0.94], and [0.33 0.41], respectively. These results indicate that crows determine their movement orientation by not only visual feedback of target location, but also the feedforward mechanism of predicted future target location in the case of fast-moving targets. The model with prediction parameter *τ* larger than 0 (rather than the parameter equal to 0) had higher likelihood in both foraging model (Fig. 1b), suggesting that crows adopted prediction of future target location in their pecking behaviour. The maximum-likelihood parameters were *τ* = 0.06 and *τ* = 0.157 s in proportional navigation model and pure pursuit model, respectively.

**Figure 2:**
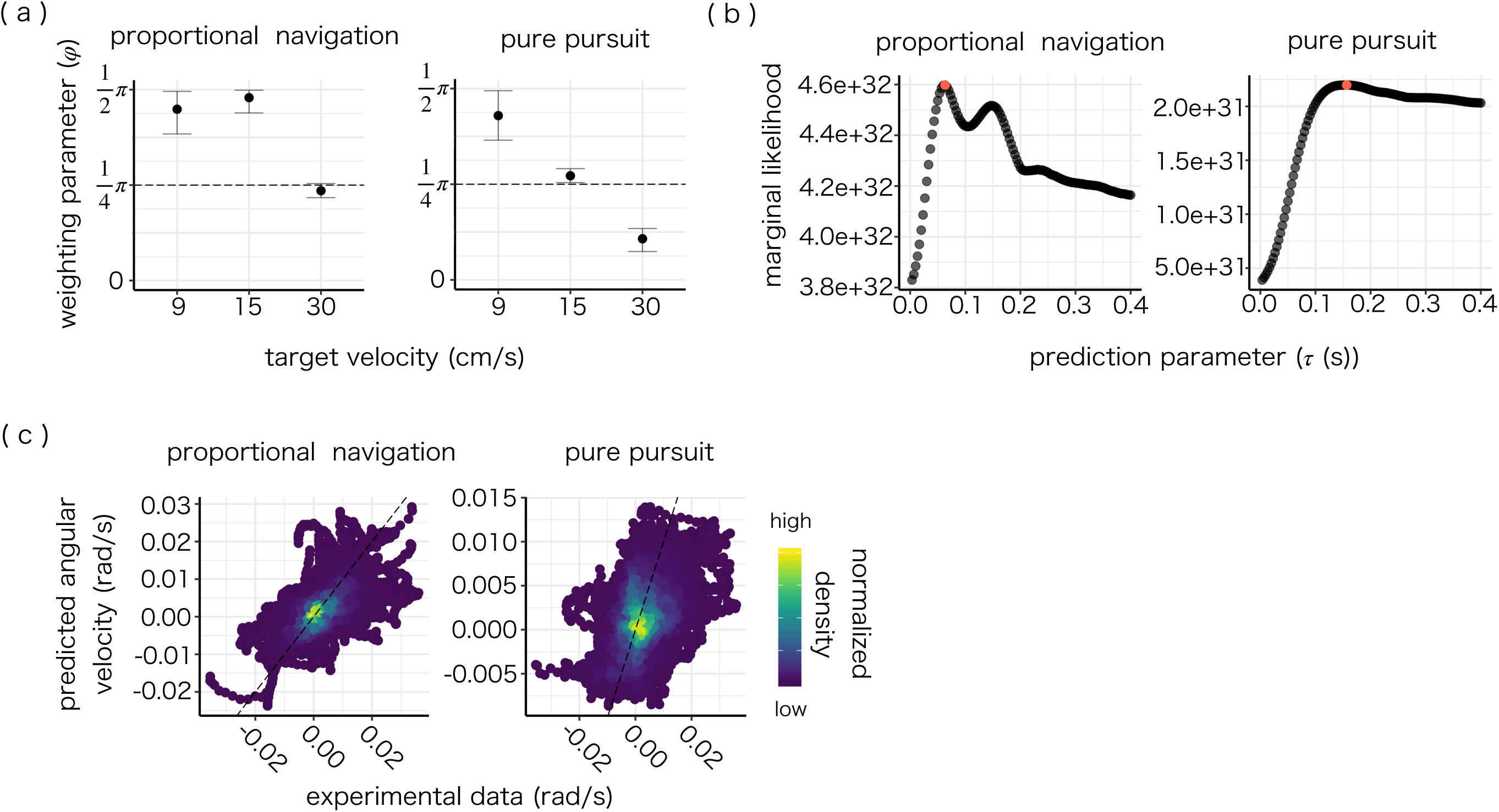
Model parameters and model prediction in each model. (a): Confidence intervals (Cis) of *φ* in each condition in proportional navigation model (left), and pure pursuit model (right). Each point denotes mean of estimated parameters. Error bars denote 95% CIs of the parameters. Dashed line represents *φ*= *π*/4 (*α* = *β* = 0.5), which indicates that feedforward predictive and visual feedback mechanisms equally contribute. (b): Calculated marginal likelihoods of models with each single *τ* in proportional navigation model (left), and pure pursuit model (right). Red points represent the highest-likelihood-*τ* in each model. (c): Heatmap representation of average of posterior predictive distribution. Estimated values are plotted against angular velocity from experimental data. Points are coloured according to the density.

The Bayes factor analysis supported the proportional navigation mechanism over pure pursuit (Bayes factor > 10,000). According to the criteria of Kass and Raftery (41), a Bayes factor value is regarded as ‘decisive’ to argue that certain model is fitted better than another, if the value is higher than 100. Our model comparison, thus, revealed that the Bayes factor was beyond ‘decisive’ criteria to argue that the proportional navigation was more suitable model. In fact, proportional navigation models have higher variance explained between real data and average of posterior predictive distribution (Fig. 2c; *R*^2^ = 0.388 in proportional navigation, and *R*^2^ = 0.172 in pure pursuit). On the basis of these results, we focused on proportional navigation rather than pure pursuit (the results of pure pursuit can be found in supplementary figures) as a plausible behavioural mechanism of interceptive movement.

Our models largely predicted the experimental data (Fig. 3a-b for well-fitted trials of proportional navigation, Fig. S1 for not well-fitted trials of proportional navigation, and Fig. S3a-b for well-fitted trials of pure pursuit). To investigate how the model fitting was altered by adding the prediction term, we subtracted the posterior distribution of the fitting, which reflect difference of fitting between with and without the prediction term. We found that the difference increased as it approached contact under the 30 cm/s condition (Fig. 3c for proportional navigation, Fig. S3c for pure pursuit). Specifically, in the successful trials, these differences became larger (Fig. 3c, red line). These results show that crows might take advantage of predictive mechanisms to generate successful foraging behaviour in the case of fast-moving targets.

**Figure 3:**
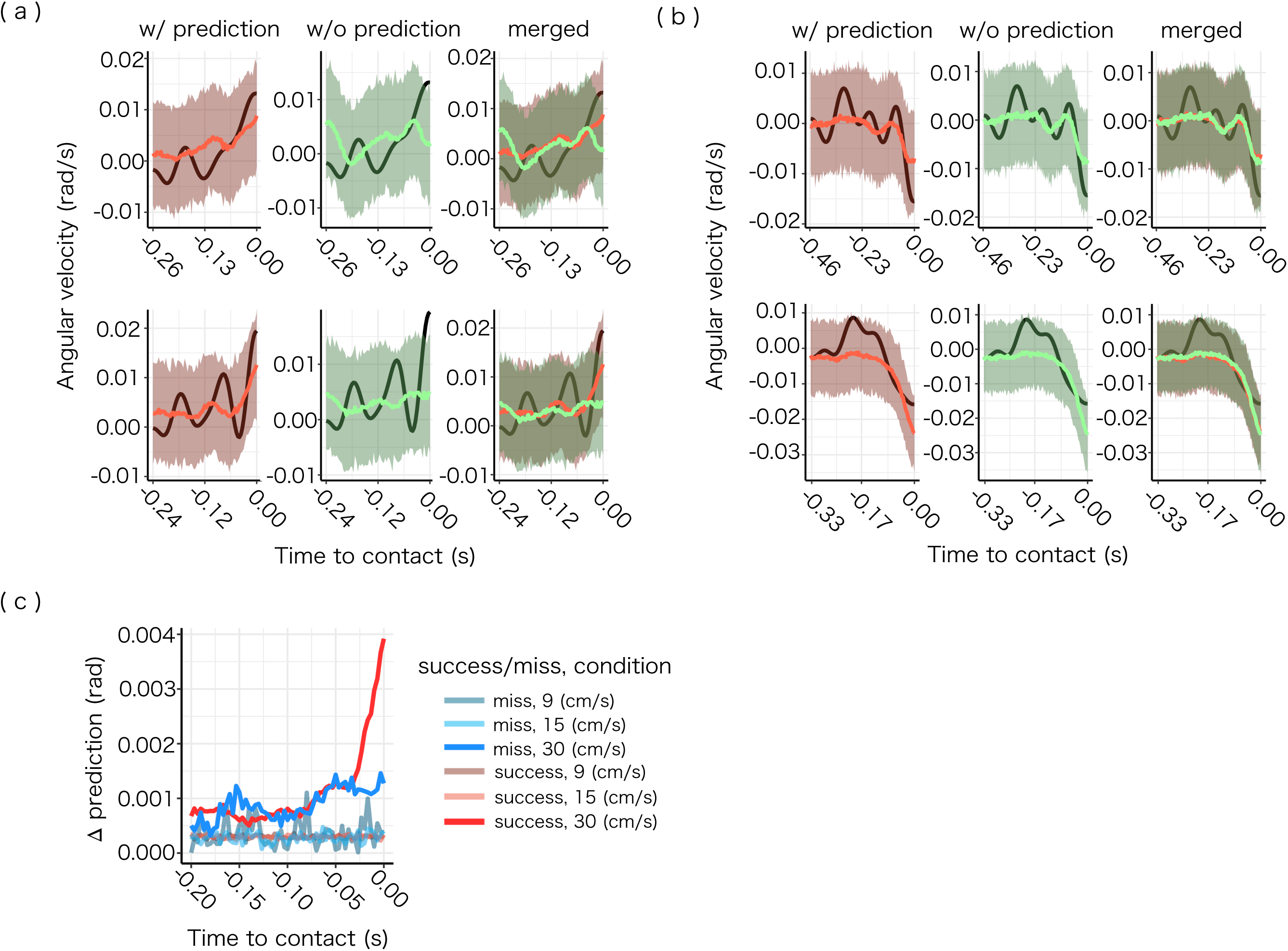
(a): Example posterior prediction intervals of two successful trials, to which the model was relatively well-fitted. Black lines represent experimental data; coloured lines represent mean of model’s predicted values; coloured shades represent 95% prediction intervals. Predictions with or without prediction term (*β*) are represented in red and green respectively. (b) Example posterior prediction intervals of two missed trials, to which the model was relatively well-fitted. (c): Differences between mean of posterior predictive distribution (coloured lines above) with and without the prediction term (*β*) in each target velocity condition of successful and missed trials. The numbers of trials (total trials (missed trials)) were 54(1), 62(4), 63(13) in target velocity conditions of 9, 15, and 30, respectively.

Under high target velocity conditions (30 cm/s), the simulated crow with a fitted parameter set could capture the target with relatively high probability (Fig. 4, proportional navigation; and Fig. S5, pure pursuit). However, the simulation predicted that successful capture does not necessarily require prediction of the future location of the target (yellow region in Fig. 4).

**Figure 4:**
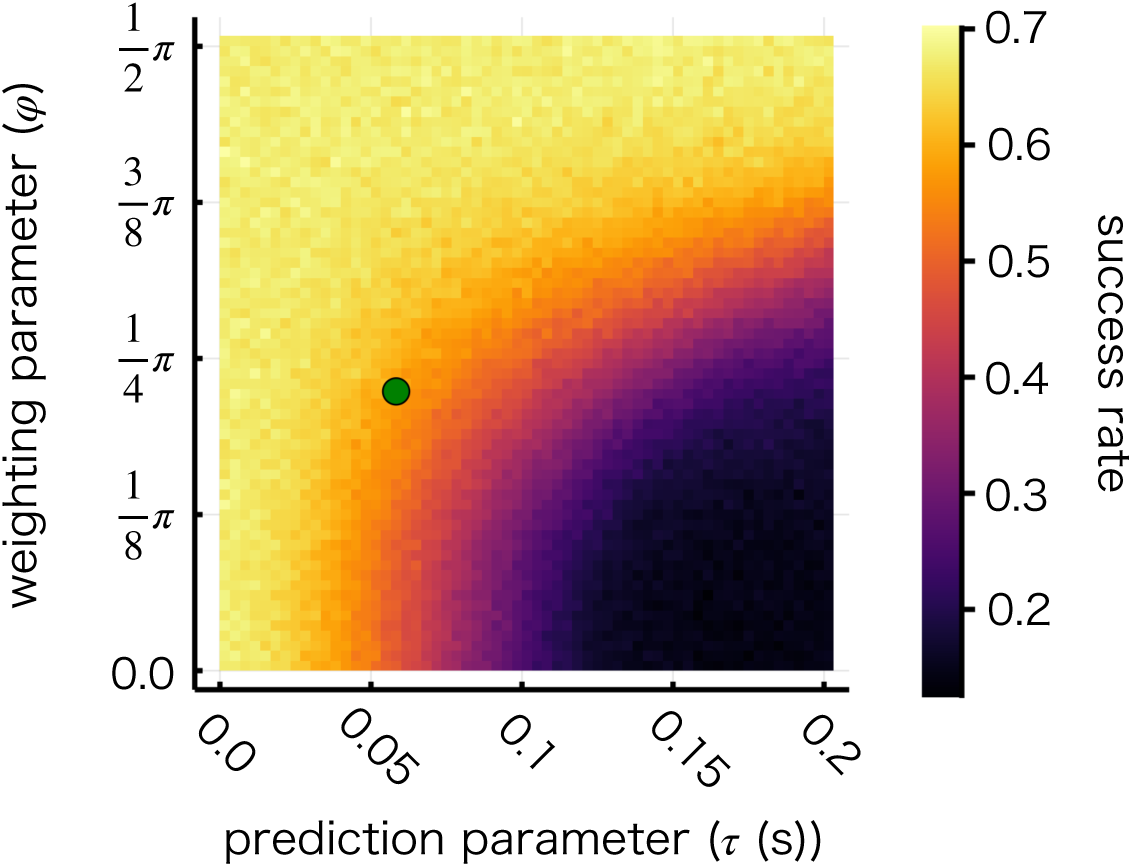
Heatmap representation of success rate in proportional navigation model with target speed 30 cm/s. Three thousand trials were simulated for each parameter set with *π/*126 step in *φ* and 1/300 step in *τ*. Green point denotes estimated parameter set to real data (the values from Fig 3a b).

**Figure 5.**
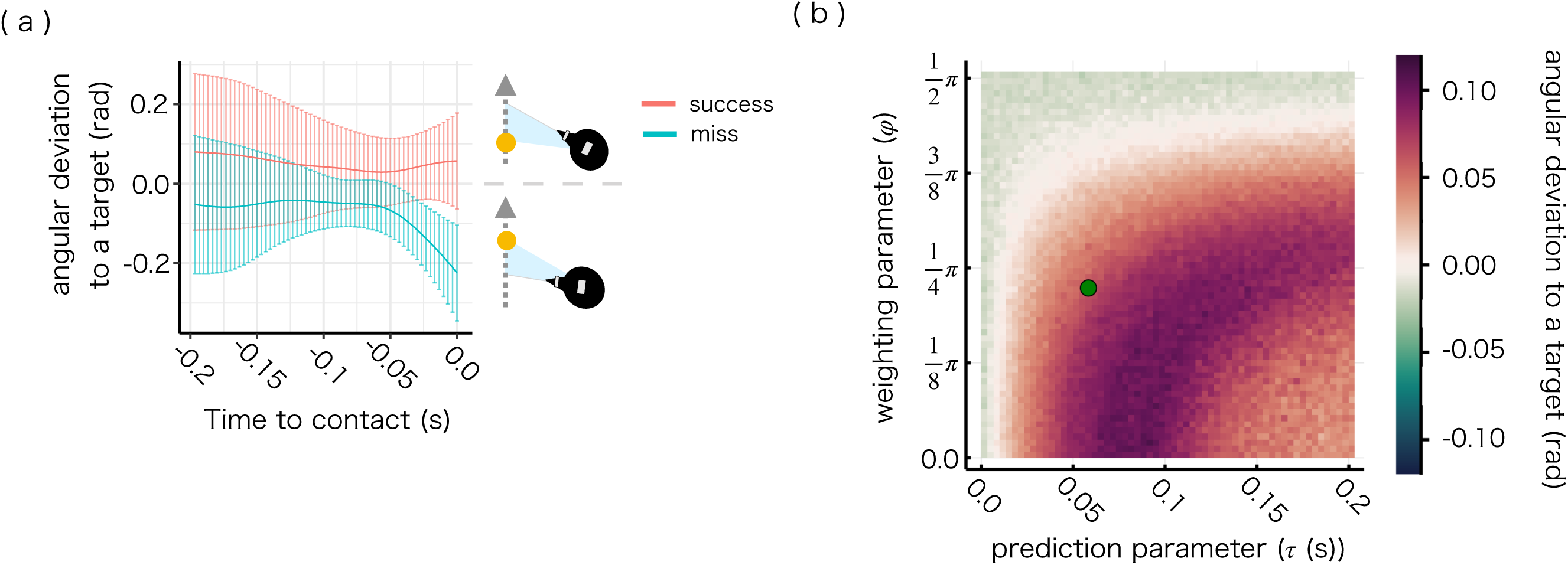
Angular deviation under high target velocity condition (30 cm/s). Positive value means crows pointed their beaks past the target direction (‘overshoot’). Negative value means crows pointed their beaks behind the target direction (‘undershoot’). (a): Experimental data. Solid lines represent mean at each time, error bars denote 1 SD. Plots are coloured according to success (red) and miss (blue). (b): Simulation data in proportional navigation model with target speed 30 (cm/s). Three thousand trials were simulated for each parameter set with *π/*126 step in *φ* and 1/300 step in *τ*. Average values of angular deviation at the time of contact in successful trials in each parameter set are plotted. Green point denotes estimated parameter set.

To test for a potential advantage of the prediction mechanism, we calculated pursuit angular deviation, which was defined as the angle between head-beak and head-target orientation (Fig. 5a). The sign of these values was determined by the orientation of the moving target: positive values indicated the crow pointed its beaks beyond the target direction; negative values indicate that the crow pointed its beak behind the target direction. It was found that the crows pointed their beaks beyond the target direction in successful trials at the time point of contact (i.e. overshooting), whereas they “undershot” the target direction in missed trials (Fig. 5a). Our simulations captured these overshooting movements in successful captures only with the prediction term (Fig. 5b, proportional navigation; Fig. S6, pure pursuit). In addition, our model with the prediction term captured quantitative features of the behaviour, in which the angular deviation was increased just before contact with the target (Fig. 5a and Fig. S2).

Under slow target velocity conditions (9 cm/s and 15 cm/s), both the fitted parameter set and the non-prediction parameter set (*β*= 0) resulted in a high success rate, with probability larger than 0.8. All parameter sets generated catching behaviour with overshooting in the successful trials as well as in the experimental data (Fig. S3). Combining success rates and pursuit angle deviation, our simulation—with the estimated parameters from real data—plausibly mimicked the interceptive behaviour of the crows.

The goals of the present study were three fold: (1) determine the behavioural mechanism governing the interceptive movements of the crows, (2) test whether the crows are using prediction of future target location, and (3) determine whether weighing of the prediction varies according to the target velocity. To resolve these questions, we compared modified versions of proportional navigation and pure pursuit models. Briefly, our computation modelling analysis supported proportional navigation as a behavioural mechanism of interceptive pecking in crows. Additionally, prediction of the future target location was adopted, especially in the interception of fast-moving targets. We describe below the mechanisms supported by the results and the implications for possible physiological implementation.

According to the comparison of models, proportional navigation is supported over the pure pursuit mechanism. Proportional navigation has been favoured in various taxa (as described in the Introduction). Specifically, predators capturing fast-moving prey tend to adopt proportional navigation. The proportional navigation mechanism is an efficient strategy in terms of computational load and is proposed to be implemented by using retinal coordinates (2, 13). To achieve proportional navigation, it is known that the animals need only to adjust the angular deviation elicited by movement of a target. In other words, proportional navigation is achieved by keeping a target in the same retinal position. This characteristic is especially advantageous in pecking, given the anatomy of birds. In birds, both the bill (i.e., effector organ) and the eyes (i.e., sensory organ) are mounted on the head, which causes an unstable and moving view associated with the pecking movement (27, 32). This difficulty does not exist in the case of primates, in which hands and arms are anatomically separated from the eyes. In birds, proportional navigation is possibly a way to overcome the difficulty derived from the unstable and moving view, because it requires only retinal coordinates of a target.

Our results are similar to those of Kane and colleagues (13), who reported that goshawks (*Accipiter gentilis*) used proportional navigation to capture evasive prey. Despite the fact that the study was focused on aerial predation, the candidate behavioural mechanism might be the same as pecking in crows, suggesting that the underlying physiological mechanisms are similar to those discussed below. Other aerial predators are also known to use proportional navigation; for example, flies dragonflies, bats, and falcons (2, 9, 10, 43). Even though our experiment focused on pecking behaviour, rather than aerial attack, our results are in agreement with these previous studies.

The simulation supports plausibility of the model and fitted parameters. The success rates indicate that fitted parameter sets were not necessarily required for successful capture. However, subsequent analysis of angular deviation revealed that fitted parameter sets mimicked the actual movement of the crows, especially under high target velocity conditions (Fig. 5b). Specifically, the crows overshot their head orientation relative to the target in successful trials, whereas they undershot in missed trials (Fig. 5a, Fig. S3a). This tendency was particularly evident when the crows shortened the distance to the fast-moving target (Fig. 5a). The model fitting was also in accordance with these results where the prediction term in the model operated in close range to the target (Fig 3c).

Predictive movements have been reported in predatory aerial steering in bats (43) and dragonflies (18) and arm-reaching movements in humans (44). In our model-based analysis, the predictive term was statistically adopted in interceptive pecking of crows. Therefore, despite the apparently different topographies of movement (aerial attack, reaching, and pecking), the predictive mechanism to capture a moving target is ubiquitous across species. As possible physiological mechanisms, it has been reported that retinal and retino-tectal circuits compute prediction of future target location. For example, there exists in birds a feedback projection from the isthmo-optic nucleus, a midbrain region, to the retina, which contributes to activation of retinal ganglion cells (45, 46) and to prediction of the future target location (47, 48). Moreover, it has been reported that retinal local circuits sensitise the adjacent ganglion cells, depending on the moving direction of the stimulus (48, 49). The possible utility of these circuits may underpin the speed of information processing, since each of these does not require pallial processing. Because predatory birds must capture fast-moving prey, it is possible that these neural circuits serve as the underlying physiological mechanism for proportional navigation with prediction.

In conclusion, the present study suggests that the interceptive pecking movement of crows is controlled by two mechanisms: prediction of future target and visual feedback. Additionally, the crows demonstrated flexibility in weighting the two mechanisms, depending on the target velocity. Proportional navigation is supported as a plausible behavioural mechanism and one that operates efficiently based on retinal coordinates. Considering the elaborate retino-tectal circuits in birds, which enable prediction of future target location, the findings of this study may provide a computational account of interceptive behaviour.

## Acknowledgments

The present research was financially supported by JSPS KAKENHI #19K21818, #19K22870, #20H01787, JST CREST #JPMJCR17A4, and Keio University Grant-in-Aid for Innovative Collaborative Research Projects #MKJ1905 to E-I. I., #16J04383 to H. M. H.M. was also financially supported by the Japan Society for the Promotion of Science, Postdoctral Fellowship for Research Abroad.

