## supplementary materials for "Interceptive capturing in large-billed crows: Velocity-dependent weighing of prediction of future target location and visual feedback of current target location"

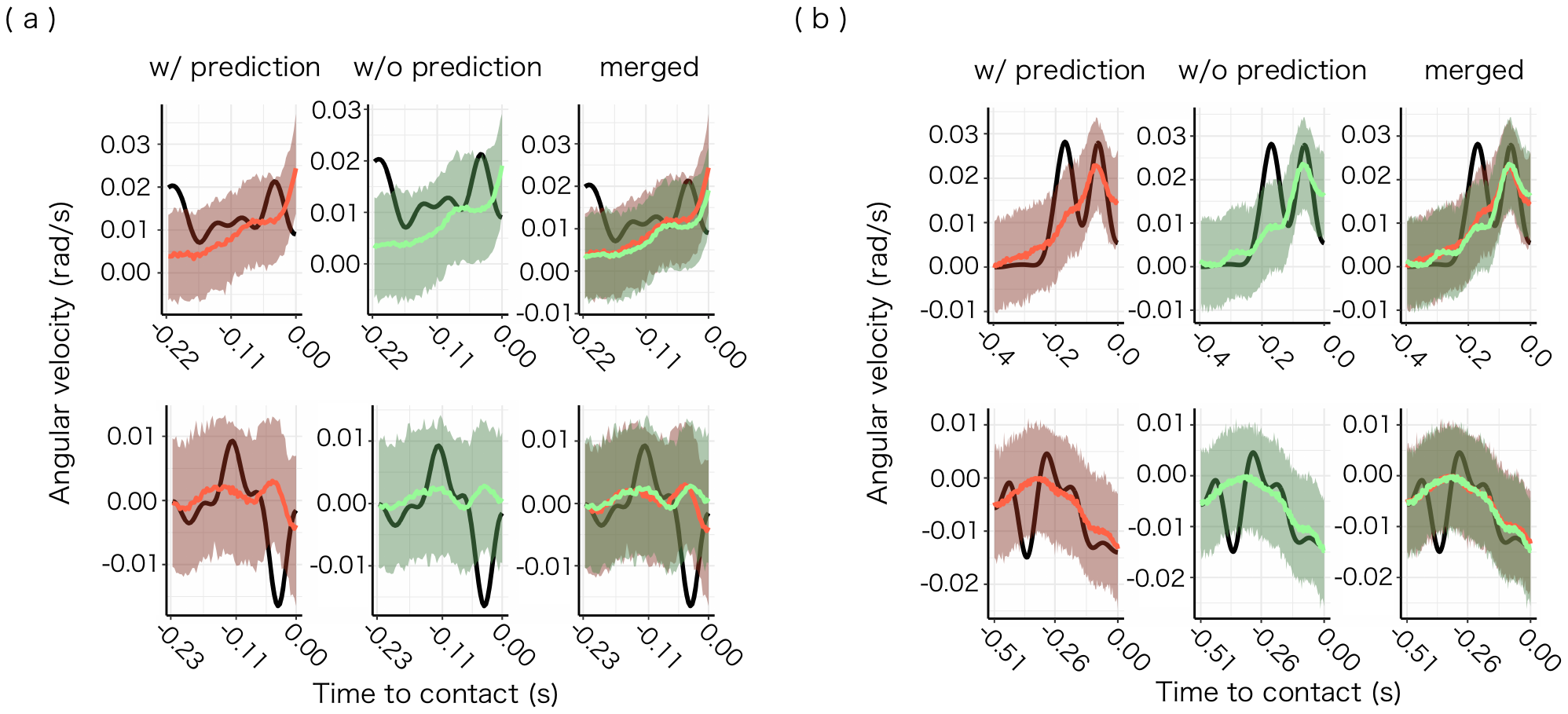

Figure S1. Example prediction intervals of proportional navigation (PN) model of successful and missed interception trials with and without the prediction term ($\beta$) and differences between models. (a): Example posterior prediction intervals of two successful trials, to which the model was not well-fitted. Black lines represent experimental data; coloured lines represent mean of predicted values in the model; coloured shades represent 95% prediction intervals. Predictions with and without the prediction term ($\beta$) are represented in red and green respectively. (b) Example posterior prediction intervals of two missed trials, to which the model was not well-fitted.

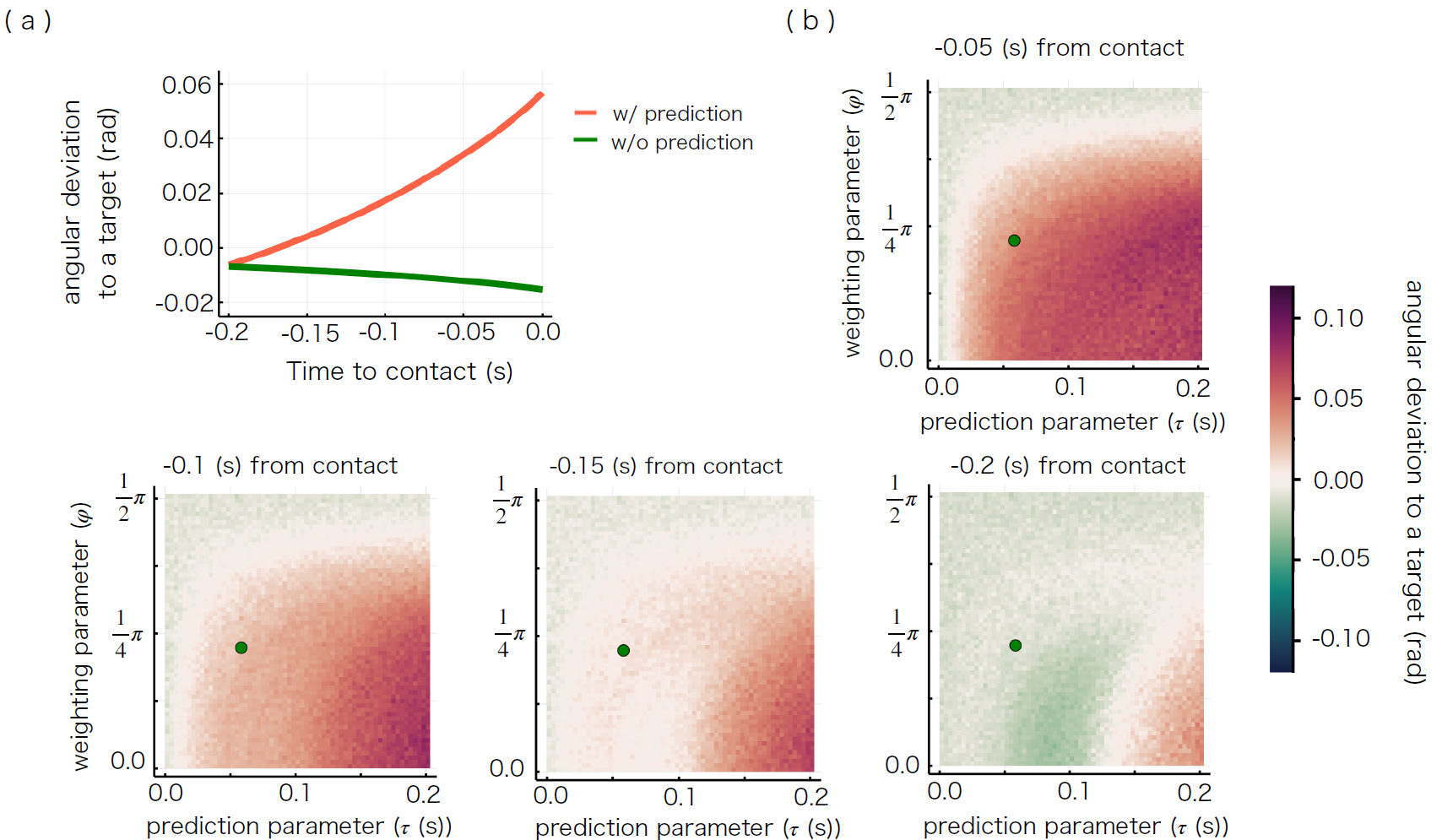

Figure S2. Time series of pursuit angle deviation under high target velocity condition (30 cm/s).

(a): Mean of the deviation at each time point. Three thousand trials were simulated in each parameter set of fitted parameters (red) and non-prediction parameter ($\tau=0,\beta=0$; green). (b): Simulation data in proportional navigation model with target speed 30 (cm/s). Three thousand trials were simulated for each parameter set with $\pi/{126}$ step in $\varphi$ and $1/{300}$ step in $\tau$. Means of pursuit angle differences at each time point (-0.05 (s), -0.1 (s), -0.15 (s), -0.2 (s) to contact in top, bottom left, bottom middle, bottom right, respectively) in successful trials in each parameter set were plotted. Green point denotes estimated parameter set.

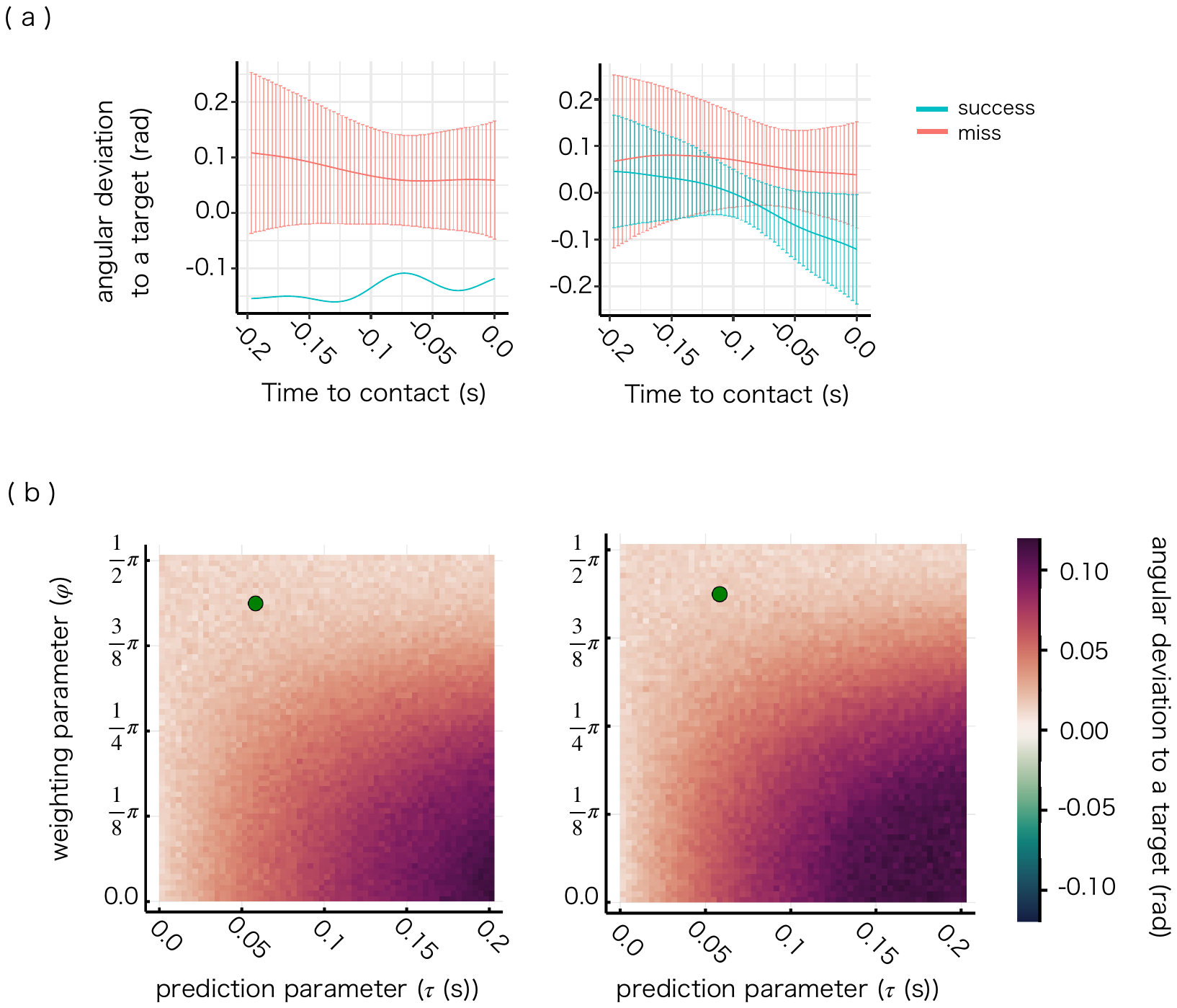

Figure S3. Pursuit angle deviation under low target velocity conditions (9 and 15 cm/s). Positive value means crows pointed their beaks past the target direction. Negative value means crows pointed their beaks behind the target direction. (a): Experimental data with target velocities of 9 cm/s (left) and 15 cm/s (right). Solid lines represent mean at each time, error bars denote SD. Plots are coloured according to successful interception (red) and missed interception (blue). (b) Simulation data in proportional navigation model with target velocities of 9 cm/s (left) and 15 cm/s (right). Three thousand trials were simulated for each parameter set with $\pi/{126}$ step in $\varphi$ and $1/{300}$ step in $\tau$. Means of pursuit angle deviation at the time of contact in success trials in each parameter set are plotted. Green point denotes estimated parameter set.

Figure S4. Example prediction intervals of pure pursuit (PP) model of successful and missed interception trials with and without prediction term ($\beta$) and differences between models. (a): Example posterior prediction intervals of two successful trials, to which the model was relatively well-fitted. Black lines represent experimental data; coloured lines represent mean of predicted values in the model; coloured shades represent 95% prediction intervals. Predictions with and without prediction term ($\beta$) are represented in red and green, respectively. (b): Example posterior prediction intervals of two missed trials, to which the model was relatively well-fitted. (c): Differences between mean of posterior distribution (coloured lines above) with and without prediction term ($\beta$) at each target velocity condition in successful and missed trials. The numbers of trials (total trials (miss trials)) were 54(1), 62(4), 63(13) at the target velocity conditions of 9, 15, and 30 cm/s, respectively.

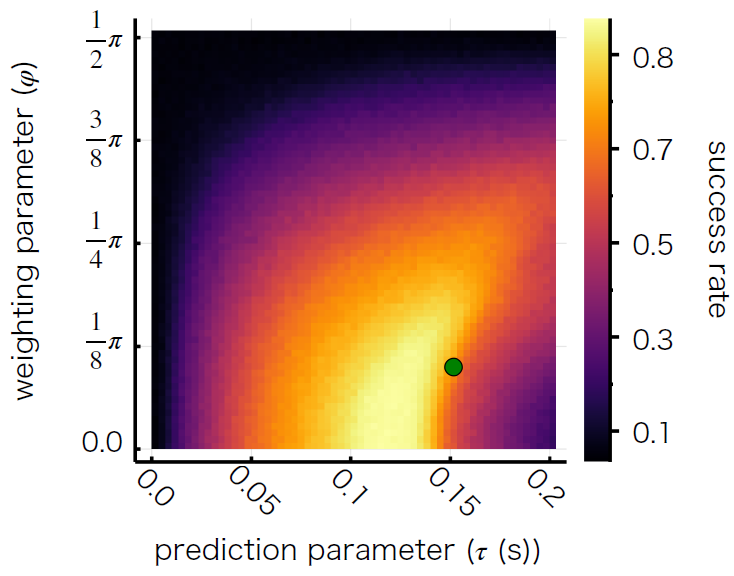

Figure S5. Success rate heat map in the pure pursuit model, with target speed of 30 cm/s. Three thousand trials were simulated for each parameter set with $\pi/{126}$ step in $\varphi$ and $1/{300}$ step in $\tau$. Green point denotes estimated parameter set.

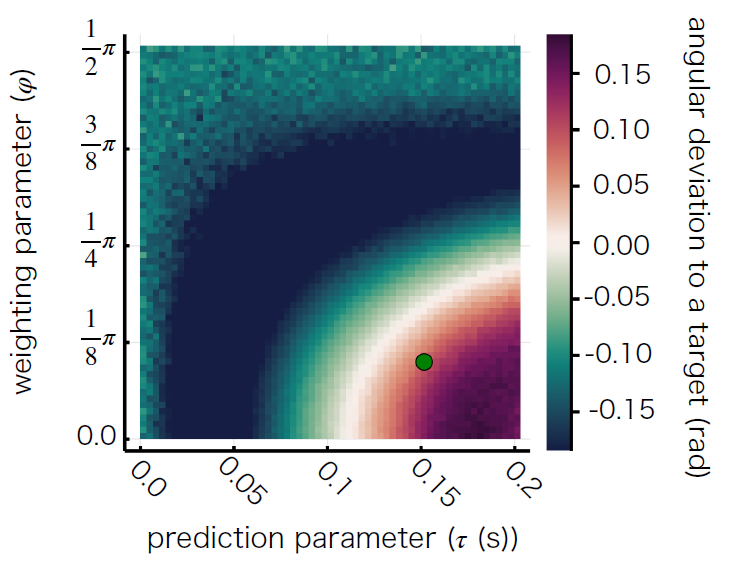

Figure S6. Angular deviation in simulation of pure pursuit model with target speed of 30 cm/s. Three thousand trials were simulated for each parameter set with $\pi/{126}$ step in $\varphi$ and $1/{300}$ step in $\tau$. Green point denotes estimated parameter set.
